# Cryopreservation of siRNA-treated cells is feasible

**DOI:** 10.1101/2025.07.01.662546

**Authors:** Melanie Sauer, Xavier Segarra-Visent, Leon Breuer, Vasileios Tzirtziganis, Tatyana Ryaykenen, David A Cooper, Dimas Echeverria, Anastasia Kremer, Reka A Haraszti

## Abstract

Cryopreservation is a routine step in the manufacturing process of adoptive cell therapies, providing critical logistic flexibility. RNAi-based therapies are increasingly being explored as enhancers or modulators of adoptive cell therapies. However, the impact of cryopreservation on cells treated with RNAi-based therapies has not been investigated before.

In this study, we addressed this knowledge gap by examining silencing efficacy in siRNA-treated cells that undergo cryopreservation. Our findings demonstrate that silencing in cryopreserved cells is comparable to that in cells maintained continuously in culture.

Moreover, we found that the duration of siRNA exposure plays a significant role in cells that later undergo cryopreservation, with extended exposure improving silencing efficiency.

However, this effect diminishes at higher siRNA concentrations.

Additionally, we showed that siRNA treatment is feasible at low temperatures (2–8°C), and siRNA-treated cells can be cryopreserved for extended periods (at least one month) without loss of efficacy. Furthermore, we demonstrated the feasibility of cryopreserving siRNA-treated primary cells, including those resembling leukapheresis material.

Our work establishes the feasibility of integrating siRNA treatments into current manufacturing processes for adoptive cell therapies.

## Introduction

RNA interference (RNAi)-based therapies are increasingly explored as modulators of adoptive cell therapies (ACT)^1-3^. The primary goals of these modulations include preventing cellular exhaustion^4^, reducing off-tumor effects^5^, modulating immune checkpoints^6,7^, enhancing efficacy^8^, and engineering desired cell phenotypes^5^.

In the manufacturing of adoptive cell therapies, the workflow often includes one or two cryopreservation steps. Hospitals with apheresis units typically harvest cells from patients. These cells are then cryopreserved and sent to a regional center, where cells are thawed, transduced with a viral vector encoding the chimeric antigen receptor (CAR) or engineered T cell receptor (TCR)^9,10^. Cells are then cryopreserved a second time and finally sent back to the treatment center^9^. Hematopoietic stem cell transplants (HSCT), the earliest form of adoptive cell therapy, are usually administered fresh within 72 hours of apheresis, but may be cryopreserved when immediate transport is not feasible^11^, when stem cell donors are unavailable at the required time^11^, or when the stem cell graft needs to be secured before starting the conditioning of the patient for HSCT^12,13^. Donor lymphocyte infusions (DLI) are a form of adoptive cell therapy intended to boost the graft-versus-leukemia effect after HSCT. DLIs are nearly always cryopreserved immediately after collection^14^, as they are often administered in a dose-escalated manner over separate sessions. These cryopreserved cells are administered to patients immediately upon thawing, except in certain cases—such as small children—where dimethyl sulfoxide (DMSO) is removed post-thaw to improve tolerance^15^. Tumor infiltration lymphocyte (TIL) manufacturing also typically involves a cryopreservation step^16^.

Introducing small interfering RNAs (siRNAs) into this process requires precise timing to maximize therapeutic impact. Due to the rapid bedside thawing of cells just before administration to patients, siRNA delivery at this stage is not feasble. Consequently, any siRNA treatment would need to occur before cryopreservation. We have recently shown, that synthetic siRNAs and miRNAs have a biologically meaningful duration of effect in rapidly dividing cells to be modulators of adoptive cell therapies^17^. However, the effect of cryopreservation on fully chemically modified siRNA efficacy has not been studied before. Optimal timing and duration for siRNA exposure prior to cryopreservation needs to be determined to ensure efficacy in modulating cellular functions.

## Results

We used a previously established fully chemically modified siRNA platform, characterized by a combination of 2’-fluoro and 2’-O-methyl modifications, along with phosphorothioate linkages^18^. This platform featured a 5’-phosphate on the guide strand and an asymmetric design, comprising a 21-nucleotide-long guide strand and a 16-nucleotide-long passenger strand, with the latter covalently conjugated to cholesterol^18^.

### Optimization of siRNA Exposure Duration Prior to Cryopreservation

To identify optimal siRNA incubation time prior to cryopreservation, we used the Jurkat suspension cell line, chosen for its reduced cell loss during freeze-thaw processes compared to adherent cells. Jurkat cells were co-incubated with siRNA for different durations (1–5 days), after which the siRNA was removed. Cells were either cultured continuously or cryopreserved for 24 hours before thawing and returning to culture.

Longer exposure to siRNA led to more robust gene silencing in cryopreserved cells, with statistically significant improvements observed as incubation time increased (Figure 1B). Specifically, a 5-day siRNA exposure resulted in the most effective knockdown (p = 0.03), followed by 4- and 3-day exposures, which were not significantly different from each other (p = n.s.) but outperformed the 2-day condition (p = 0.0042). One-day exposure showed the least silencing efficacy (p = 0.062). In contrast, these differences were not statistically significant in continuously cultured cells (Figure 1A).

**Figure 1.**
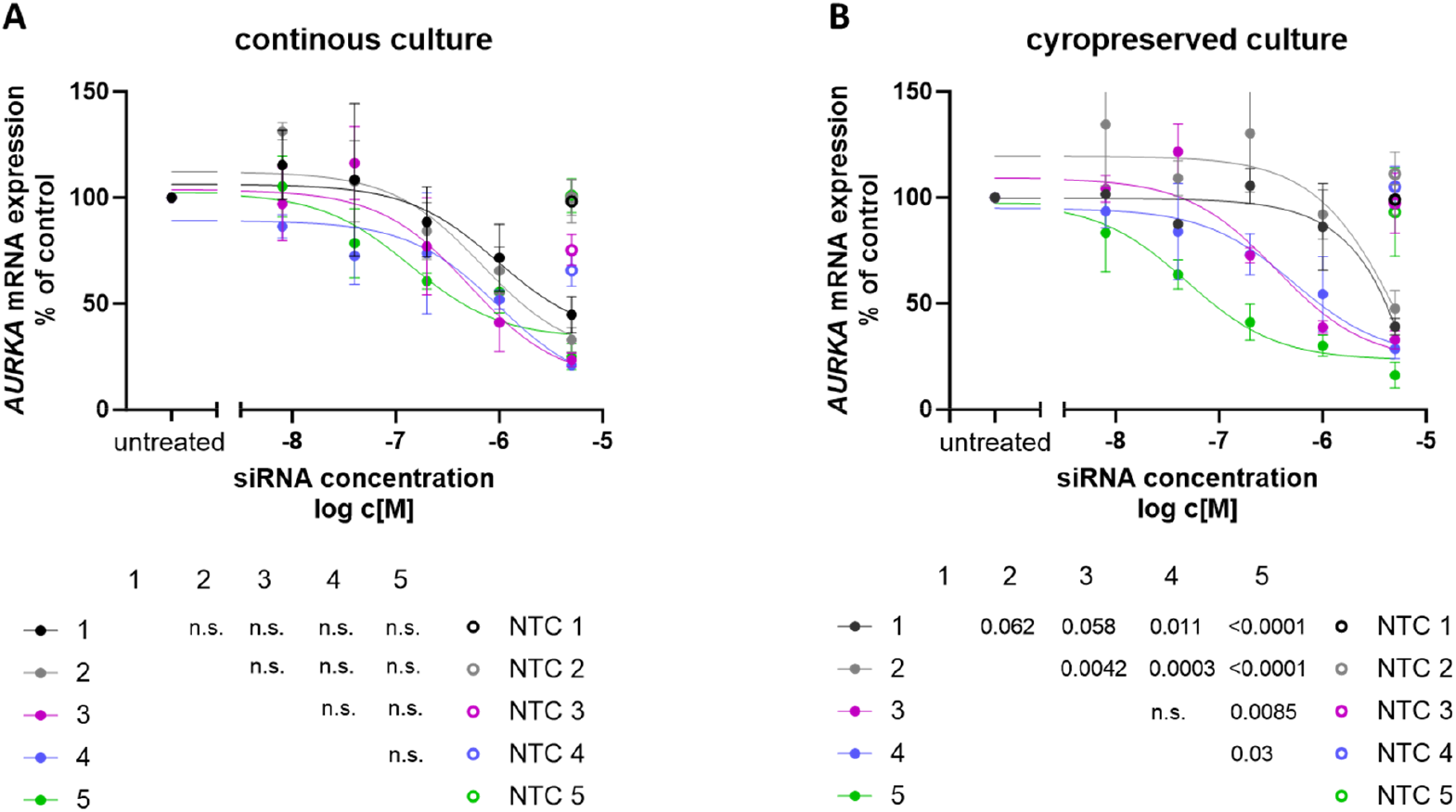
Effect of siRNA Exposure Duration to silencing efficacy. Jurkat cells were co-incubated with siRNA targeting *AURKA* at various concentrations, as indicated on the x-axis, and for different durations, represented by the color code. After the specified exposure time to the siRNAs, the medium containing the siRNA was removed by centrifugation and replaced with fresh medium. The cells were then either continuously cultured further in an incubator (A) or cryopreserved at –80 °C for 24 hours (B). Following the freezing period, the cells were thawed, the freezing medium was removed, fresh medium was added, and the cells were further cultured in an incubator. All cells were cultured for a total of 6 days. Subsequently, cells were lysed and mRNA levels were quantified using the QuantiGene Singleplex assay. mRNA expression was normalized to the housekeeping gene and untreated control. mean ± SEM, N = 4.

Importantly, at high siRNA concentrations (5 µM), these differences disappeared, indicating that maximal target engagement can overcome timing-dependent variability. Based on these findings, we selected a 24-hour co-incubation with high-concentration siRNA (5 µM) as the most practical and robust condition for downstream experiments, given its compatibility with clinical manufacturing timelines.

Interestingly, cryopreserved cells showed slightly better silencing than their continuously cultured counterparts at longer exposure durations (4 days: p = 0.06; 5 days: p = 0.01; paired t-test).

### Effect of Cryopreservation Duration on Silencing Efficacy

We next assessed whether the length of time cells were stored cryogenically would influence silencing. Cells exposed to siRNA and then cryopreserved for either 24 hours or one month showed comparable knockdown efficiencies to those cultured continuously (Figures 2A and 2B), indicating that the duration of cryopreservation does not adversely affect the stability or function of siRNA-loaded cells. These findings were independently validated in HeLa cells using a different siRNA sequence targeting the *RAN* gene, confirming the generalizability of our approach (Supplementary Figure 1).

**Figure 2.**
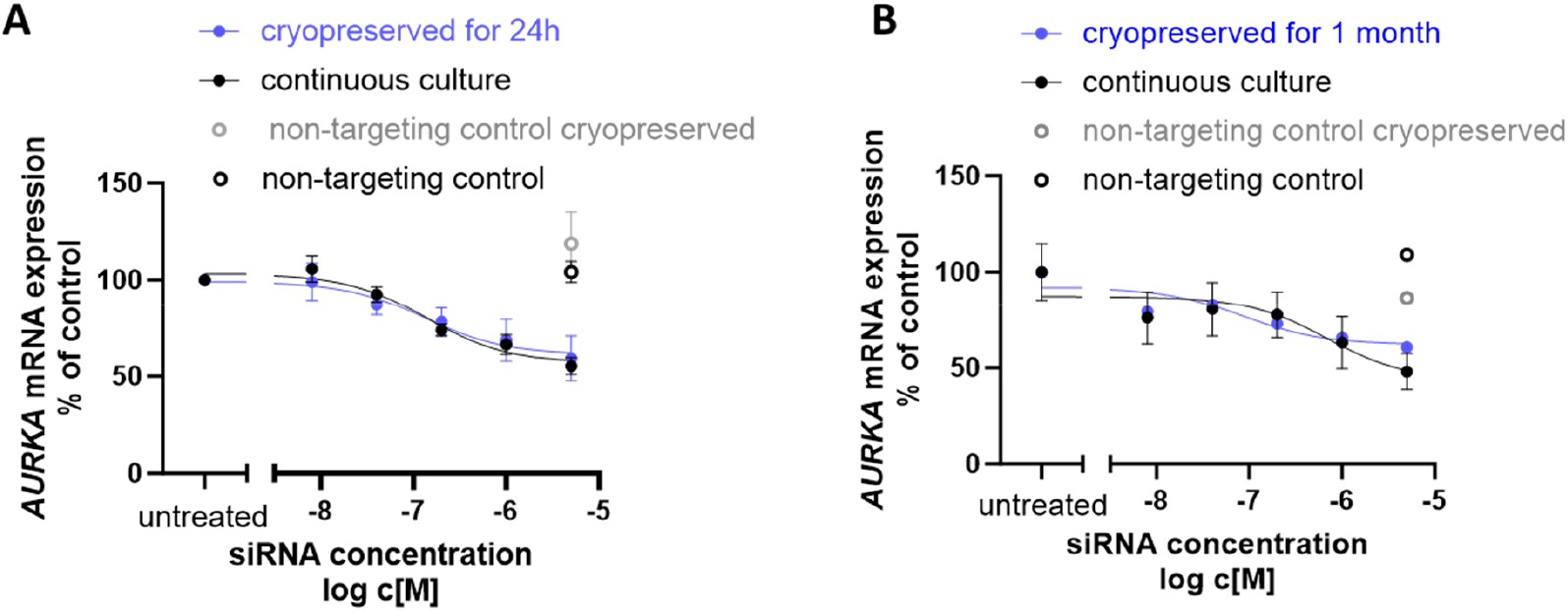
Cryopreservation of siRNA-treated cells maintains silencing efficacy. Jurkat cells were exposed to siRNA targeting *AURKA* at various concentrations, as indicated on the x-axis for 24 hours. siRNA-containing medium was then removed and cells cryopreserved for either 24 hours (A) or 1 month (B) at –80 °C. Cell were then thawed and cultured further in an incubator for 5 days. Subsequently, cells were lysed and mRNA levels were quantified using the QuantiGene Singleplex assay. mRNA expression was normalized to the housekeeping gene and untreated control. mean ± SEM, N = 3-6.

**Figure 3.**
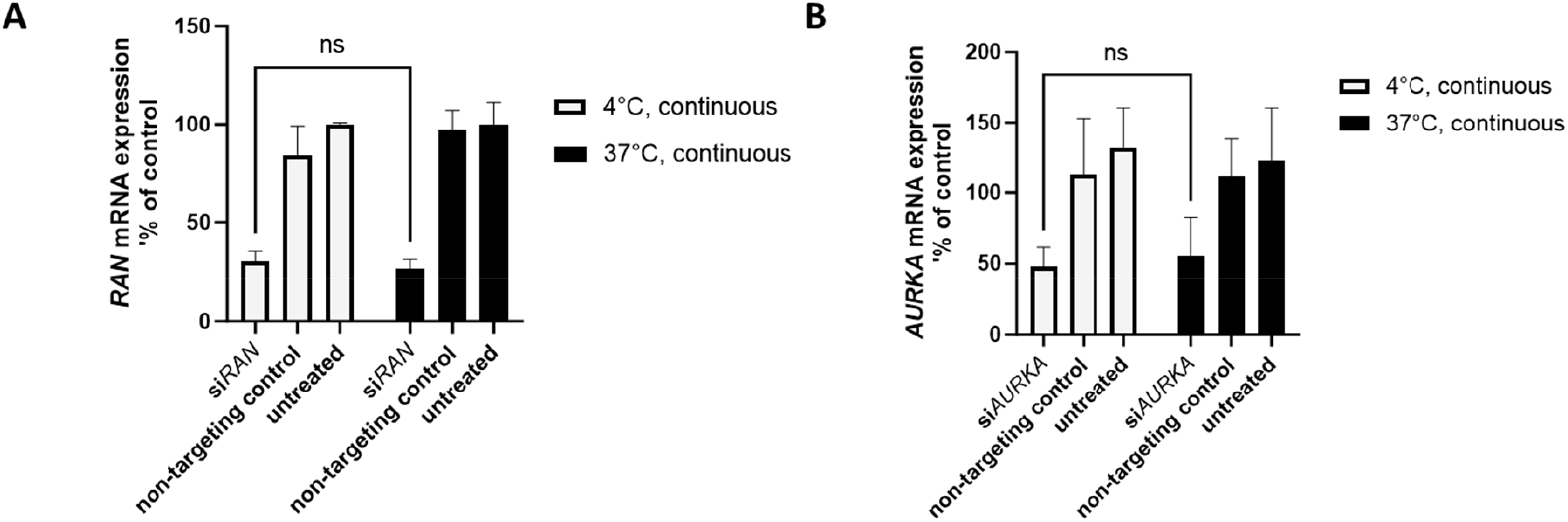
Productive siRNA uptake is maintained at 4°C. Peripheral blood mononuclear cells (PBMCs) were treated with siRNA targeting *RAN* (A) or *AURKA* (B) at either 4°C (empty bars) or 37°C (black bars) for 24 hours. siRNA-containing medium was then removed and replaced with fresh medium. All cells were now placed into a 37°C cell culture incubated and cultivated further for 5 days. Subsequently, cells were lysed and mRNA levels were quantified using the QuantiGene Singleplex assay. mRNA expression was normalized to the housekeeping gene and untreated control. mean ± SEM, N = 3-6.

### Impact of Temperature During siRNA Exposure

Given that apheresis-derived cells are typically stored at 4°C prior to processing and cryopreservation, we tested whether siRNA treatment could be effective at this lower temperature. Primary human peripheral blood mononuclear cells (PBMCs)—a clinically relevant heterogeneous population similar to those used in adoptive cell therapies such as CAR T or DLI—were treated with two independent siRNA sequences at either 37°C or 4°C for 24 hours. In both cases, comparable levels of gene silencing were observed, indicating that siRNA delivery is feasible under cold storage conditions. This finding significantly broadens the translational applicability of our approach to real-world clinical workflows.

## Discussion

Cryopreservation plays a critical role in the logistics and scheduling flexibility of adoptive cell therapies, enabling decoupling of manufacturing steps and facilitating global distribution. This importance was notably underscored during the COVID-19 pandemic, during which cryopreservation ensured the continuity of hematopoietic stem cell transplantation (HSCT) despite widespread logistical disruptions^12,13^. Moreover, cryopreservation is now an integral component of the manufacturing process for all approved CAR-T and engineered TCR-T cell therapies^19^.

As RNAi therapeutics are increasingly explored as modulators or potentiators of therapeutic immunity and adoptive cell therapies, we sought to address a knowledge gap regarding the feasibility of cryopreserving siRNA-treated cells. Our study demonstrates that cryopreservation of siRNA-treated cells, including both cell lines and primary cells resembling human apheresis material, is feasible. siRNA silencing efficacy in cryopreserved cells was found to be comparable to that observed in continuously cultured cells. Interestingly, we observed that the duration of siRNA exposure had a greater impact on silencing efficacy in cryopreserved cells than in continuously cultured cells. This phenomenon may be explained by several factors. Cryopreservation has been shown to synchronize cellular circadian rhythms^20^, which may, in turn, influence RNAi machinery components^21^, cellular uptake pathways^22,23^, or endosomal trafficking^24,25^. Additionally, stress responses induced during freezing and thawing, along with delayed post-thaw recovery, may further modulate gene silencing outcomes^19^. We also show that siRNA-treated cells can be stored long-term in cryogenic conditions and, upon thawing, retain their silencing potency.

Notably, siRNA can be taken up to cells and support efficient silencing at cold temperatures (2–8°C) – a condition relevant for adoptive cell therapy handling and storage before cryopreservation. This is a surprising finding, given the generally inefficient nature of endocytosis at low temperatures^26,27^. However, this phenomenon is consistent with previous reports showing that liposomal uptake can proceed effectively under similarly cold conditions^28^.

Overall, our findings provide strong evidence that treating cells with siRNA drugs at cold temperatures and cryopreserving siRNA-treated cells is a viable approach without compromising silencing efficacy. These results support the integration of siRNA-based interventions at early stages of adoptive cell therapy manufacturing – specifically, treating fresh leukapheresis material before cryopreservation – thereby enabling broader use of RNAi to fine-tune therapeutic cell functions without introducing additional complexity into established workflows.

## Methods

### Oligonucleotides

Compounds were synthesized using standard solid-phase phosphoramidite chemistry on either a Dr. Oligo 48 high-throughput RNA synthesizer (Biolytic) or using a MerMade 12 (BioAutomation) synthesizer. Standard RNA 2′-*O*-methyl and 2′-fluoro modifications were applied for improving siRNA stability (Chemgenes). The sense strands were synthesized at a 1-μmol scale on a cholesterol-functionalized controlled pore glass (CPG) solid support (Chemgenes) for in vitro experiments. 5-μmol scale was used for in vivo experiments. For post-synthesis deprotection, sense strands were cleaved from the CPG and deprotected using 40% aqueous methylamine and 30% NH_4_OH (1:1, v/v) at room temperature for 2 h. Guide strands were cleaved and deprotected with 30% NH_4_OH containing 3% diethylamine for 20 h at 35°C. 5′-E-VP containing antisense strands were washed with bromotrimethylsilane:pyridine (3:2, v/v) in dichloromethane while still on solid support previous to deprotection. The deprotected oligonucleotide solutions were filtered to remove CPG residues and dried under vacuum. Compounds were precipitated using a modified ethanol precipitation protocol. The oligonucleotides were quality-controlled by liquid chromatography-mass spectrometry (LC-MS). Desalting was carried out by size exclusion chromatography.

### Cell culture

Jurkat cells were cultured in RPMI-1640 medium containing stabilized glutamine and sodium bicarbonate (R2405, SIGMA), supplemented with 10% fetal bovine serum (FBS) (11573397, Fisher Scientific), 1% Penicillin-Streptomycin (P/S) (P0781, SIGMA), and 25 mM HEPES buffer (9157.1, Carl Roth).

HeLa cells were maintained in the same RPMI-1640 base medium with stabilized glutamine and sodium bicarbonate, supplemented with 10% FBS and 1% P/S.

Peripheral blood mononuclear cells (PBMCs) were isolated via density gradient centrifugation. The buffy coat was first decontaminated using 70% ethanol, transferred into sterile conical tubes, and diluted at a 1:1 ratio with phosphate-buffered saline (PBS). This mixture was gently layered over Ficoll (11768538, Fisher Scientific) and centrifuged to separate mononuclear cells. The interphase containing PBMCs was carefully extracted, washed with PBS, and centrifuged again to eliminate residual platelets. The resulting cell pellet was resuspended in RPMI-1640 medium containing 10% FBS (11573397, Gibco), 1% P/S (P0781, Sigma), 25 mM HEPES (9157.1, Carl Roth) and 1 mM sodium pyruvate. Cells were subsequently counted using a Neubauer hemocytometer.

Hela cell experiments were carried out in 96-well flat bottom plates, experiment with Jurkat cells and PBMCs were carried out in 96-well U-bottom plates.

### Cryopreservation

PBMCs, Jurkat and HeLa cells were centrifuged at 300 g for 7 minutes and the supernatant discarded. Cell pellet was resuspended in freezing medium (90% FBS (11573397, Fisher Scientific) with 10% DMSO (1198621, Omnilab) for PBMCs and 9% FBS, 10% DMSO in RPMI) and transferred to -80°C. The cells were then centrifuged at 300 g for 7 minutes and the DMSO-containing medium was fully removed. The cell pellet was resuspended in new medium and transferred to a 96-well plate for further incubation.

### mRNA quantification

mRNA levels were measured using the QuantiGene Singleplex assay. A fresh Working Lysis Mixture was prepared before each use by diluting Proteinase K (QS0106, Life Technologies GmbH) into the Lysis Mixture at a 1:100 ratio. This Working Lysis Mixture was then added to cell samples at a 1:2 ratio (v/v). Samples were mixed thoroughly and incubated at 55°C for 30 minutes. After incubation, the lysates were mixed again by pipetting and either immediately processed or stored at −80°C for later analysis. Prior to further processing, frozen samples were fully thawed at room temperature and then incubated at 37°C for 15 minutes.

The QuantiGene Singleplex (QGS) assay was performed following the manufacturer’s instructions, using the Invitrogen™ QuantiGene™ Sample Processing Kit for cultured cells (QS0103, Life Technologies GmbH) and the QuantiGene™ Singleplex Assay Kit (QS0016, Life Technologies GmbH). Hypoxanthine-guanine phosphoribosyltransferase (HPRT) served as the reference housekeeping gene. The specific probe sets used in the assay were: AURKA (SA-10135), GAPDH (SA-10001), HPRT (SA-10030), and RAN (SA-15837).

### Data analysis and visualization

Data analysis and visualization were conducted using GraphPad Prism (Version 10.1.1 (323)). Silencing curves were fitted using the “log(inhibitor) vs. response (three parameters)” function. Comparisons of these curves were performed using two-way ANOVA with multiple comparison correction or paired t tests. Shapiro-Wilk normality tests confirmed normal distribution of data.

## Acknowledgement

We thank Anastasia Khvorova for making oligonucleotide synthesis infrastructure available for this project. This work was supported by the German Cancer Aid [70113948 to R.A.H.]; and the Faculty of Medicine, University of Tübingen [473-0-0 to R.A.H., 2652-0-0 to R.A.H.]. R.A.H. was further supported by the Deutsche Gesellschaft für Innere Medizin Clinican Scientist Program and by the MINT-Clinician Scientist Program of the Medical Faculty Tübingen, funded by the Deutsche Forschungsgemeinschaft (493665037).

## Author Contributions

Conceptualization R.A.H. Methodology. X.S., A.K., T.R., D.E. Investigation M.S., X.S., L.B., V.T., D.A.C. Writing – Original Draft R.A.H. Visualization M.S., X.S., L.B., VT., R.A.H. Supervision R.A.H. Project Administration R.A.H. Funding Acquisition R.A.H.

---

**Supplementary Figure 1.**
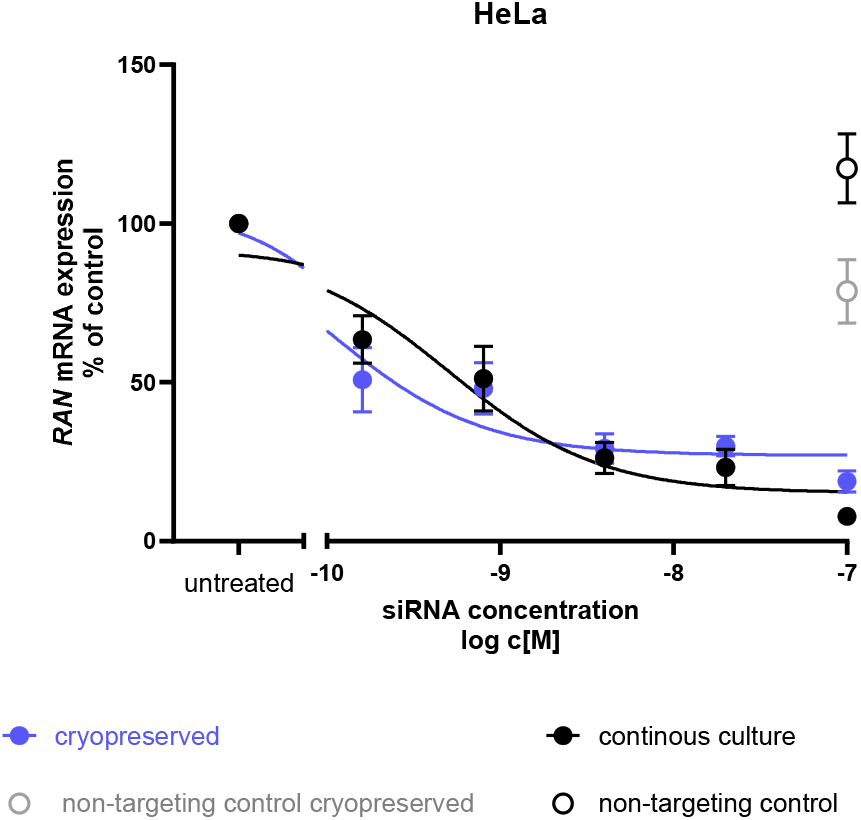
Cryopreservation of siRNA-treated cells maintains silencing efficacy in adherent cells. HeLa cells were treated with siRNA targeting AURKA at varying concentrations, as indicated on the x-axis, for 24 hours. Following exposure, the siRNA-containing medium was removed, and the cells were cryopreserved at –80 °C for 24 hours. After thawing, cells were transferred to fresh medium and cultured for an additional 5 days under standard incubator conditions. Subsequently, cells were lysed and mRNA expression was analyzed using the QuantiGene Singleplex assay. Expression levels were normalized to both a housekeeping gene and the untreated control. mean ± SEM, N = 9.

**Supplementary Table 1.**
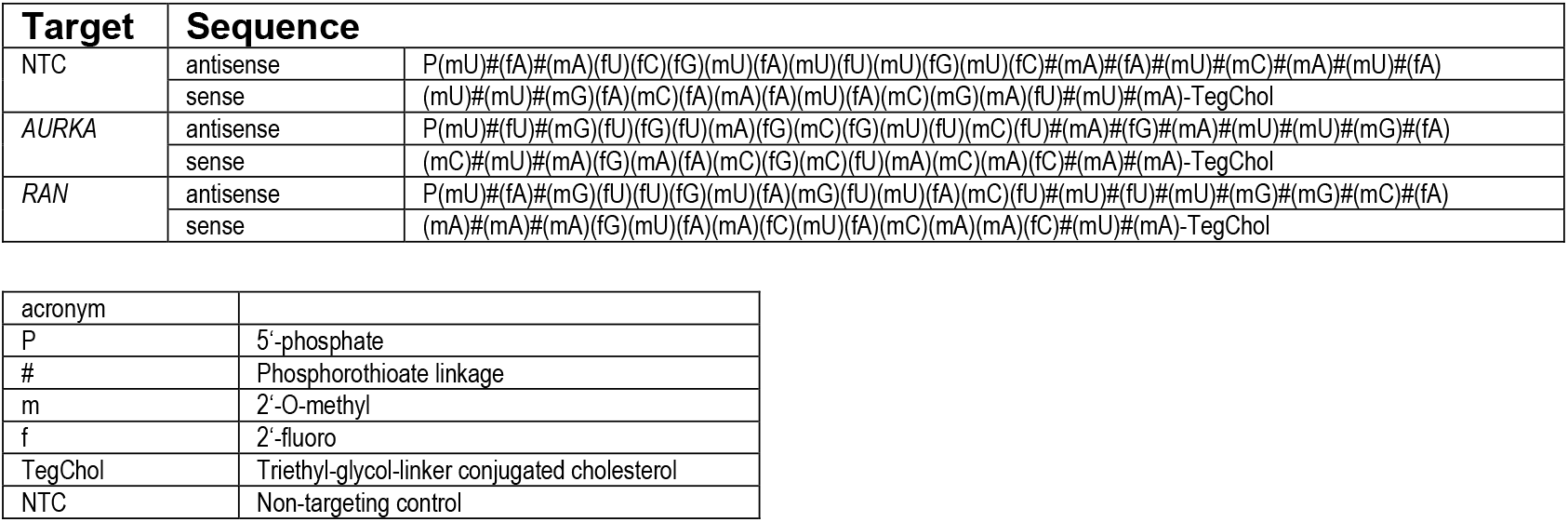
Oligonucleotide sequences and their chemical modifications used in this study

